# Dynamics of bacterial cell proliferation inhibited by specific proteins for effective horizontal transmission of the integrative and conjugative element ICE*clc*

**DOI:** 10.1101/512285

**Authors:** Sotaro Takano, Akiko Koto, Ryo Miyazaki

**Affiliations:** Bioproduction Research Institute, National Institute of Advanced Industrial Science and Technology (AIST), Tsukuba, Japan; Computational Bio Big Data Open Innovation Laboratory (CBBD-OIL), AIST, Tsukuba, Japan; Faculty of Life and Environmental Sciences, University of Tsukuba, Tsukuba, Japan

## Abstract

The integrative and conjugative element ICE*clc*, a prevalent mobile element in proteobacteria, is one of the experimental models for horizontal gene transfer. ICE*clc* is usually retained in a bacterial chromosome, but can be excised and transferred from the donor to other bacterial lineages. The horizontal transmission is accomplished by developing specialized transfer competent (tc) cells in the donor population. The tc cells entirely dedicate to the ICE transmission by sacrificing their proliferation, which results in an increase in the transfer frequency. The cell growth impairment is mediated by two specific genes, *parA* and *shi*, on ICE*clc*, but its mechanistic details and cellular dynamics are still unknown. We here developed fluorescence reporter strains to monitor intracellular behavior of ParA and Shi proteins as well as host cellular proliferation at the single-cell level. Superresolution imaging revealed that ParA colocalized with the host nucleoid while Shi diffused in cytoplasm during the growth impairment. Mutations in the Walker A motif of ParA diminished the inhibitory effect. Combining quantitative time-lapse microscopy and numerical simulations using mathematical models, we found that ParA and Shi initially blocked cell division and then, as time elapsed, inhibited cellular elongation. The *parA-shi* locus is highly conserved in other ICEs, and the ParA-Shi-mediated growth inhibition was still observed in different proteobacterial species, suggesting that the ICEs have evolved the system to efficiently distribute themselves in the niche. The results of our study provide mechanistic insight into the novel and unique system on ICEs and help to understand such epistatic interaction between ICE genes and host physiology that entails efficient horizontal gene transfer.

**Author summary:** Horizontal gene transfer is a major diving force for bacterial evolution, which is frequently mediated by mobile DNA vectors, such as plasmids and bacteriophages. Integrative and conjugative elements (ICEs) are a relatively new class of mobile vectors, which normally integrate in a host chromosome but under certain conditions can be excised and transferred from the host to a new recipient cell via conjugation. Recent genomic studies estimated that ICEs are more abundant than plasmids among bacteria. Why so prevalent? One of the characteristics of ICE*clc*, an ICE model in proteobacteria, is that it develops specialized cells which entirely dedicate to the ICE horizontal transmission by repressing their proliferation. Here, we qualitatively and quantitatively describe two proteins, which are expressed from ICE*clc* when it transfers, and how they actually inhibit the host cell growth. Our results suggest that the system has evolved for the efficient horizontal transmission and is widely conserved in the ICE family.

## Introduction

Horizontal gene transfer is a pivotal event for prokaryotic evolution, since large pieces of DNA can be exchanged among bacterial species. Horizontal gene transfer is frequently, but not exclusively, mediated by mobile DNA vectors such as conjugative plasmids and bacteriophages. Integrative and conjugative elements (ICEs) are another class of ubiquitous mobile vectors that are usually retained in bacterial chromosomes but under certain conditions can be excised and transferred from the donor to other bacterial lineages where they again integrate into the new host chromosomes [1–3]. A large number of ICEs have been experimentally identified in very different species of both gram-positive and gram-negative bacteria [4], and many more potential ICEs could be inferred from bioinformatic approaches [5–7]. Like other mobile vectors, ICEs often carry cargo genes that encode distinct features, such as antibiotic resistance and metabolic functions, and thus accelerate microbial evolution and adaptation [4]. It has been estimated that conjugative systems of ICEs are more abundant among bacteria than those of plasmids [5].

While the mechanistic basis for the pervasiveness of ICEs in the prokaryotic world are still poorly understood, part of it could be explained by their characteristic lifestyle. Once inserted in the host chromosome, ICEs will be faithfully copied in every dividing cell. As long as the integrated ICEs do not impose major fitness costs on the host, or even provides selective benefit by cargo genes, it will be stably maintained in the cells. Indeed, previous studies have reported very limited fitness costs (below 1%) or rather direct benefits from the integrated from of ICE on the whole population by silencing its activities [8,9]. In contrast, the process of horizontal transmission can be very costly, because ICEs must excise from the chromosome and induce the donor cells to produce DNA transfer machineries with associated factors [10,11]. The fact that horizontal transmission of ICE typically occurs at a very low rate (10^−2^ to 10^−7^ per donor cell) also suggests that it is sufficiently disadvantageous [1]. An attracted question is thus how ICEs have evolved their systems to deal with cost, efficiently distribute themselves, and increase their fitness (i.e. copy numbers) in a given ecological niche.

We study ICE*clc*, an 103-kb ICE originally found in *Pseudomonas knackmussii* B13 and widely distributed in proteobacteria [12,13], as an experimental model to understand evolution and adaptation of ICEs with host bacteria. We have shown that horizontal transmission of ICE*clc* necessitates development of the host bacterial cells into a transfer competence (tc) state, which occurred in only 3-5% of the stationary phase cells in a clonal population [14]. The development of tc cells is implemented with stochastic intracellular variability of regulatory molecules and subsequent bistable expression of ICE*clc* genes [11,15,16], although substantial events of excision and transfer predominantly occurred only when tc cells have been presented with new nutrients [17]. These suggest that ICE*clc* transfer is energetically costly for individual donor cells and thus restricted in a small subpopulation. Intriguingly, tc cells do not only commit to ICE*clc* transfer, but their proliferation is also impaired by simultaneous expression of two ICE*clc* genes, *parA* and *shi*, annotated as encoding a nucleotide partitioning ATPase and a hypothetical protein, respectively [14]. The *parA* and *shi* are located within a gene cluster adjacent to the *attL* end, one of the boundaries between the host chromosome and integrated ICE*clc* (Fig. 1A). Induction of the two genes reproduces the cellular growth impairment with abnormal cell morphologies, whereas their deletion abolishes the growth inhibition and, importantly, reduces the ICE transfer frequency [14]. The growth impairment by *parA* and *shi* is thus thought to benefit the overall transfer success of ICE*clc* by dedicating tc cells to the mission and increasing the chance to contact recipients [14,17,18]. However, mechanistic details and cellular dynamics of the inhibition remain unknown.

**Fig 1.**
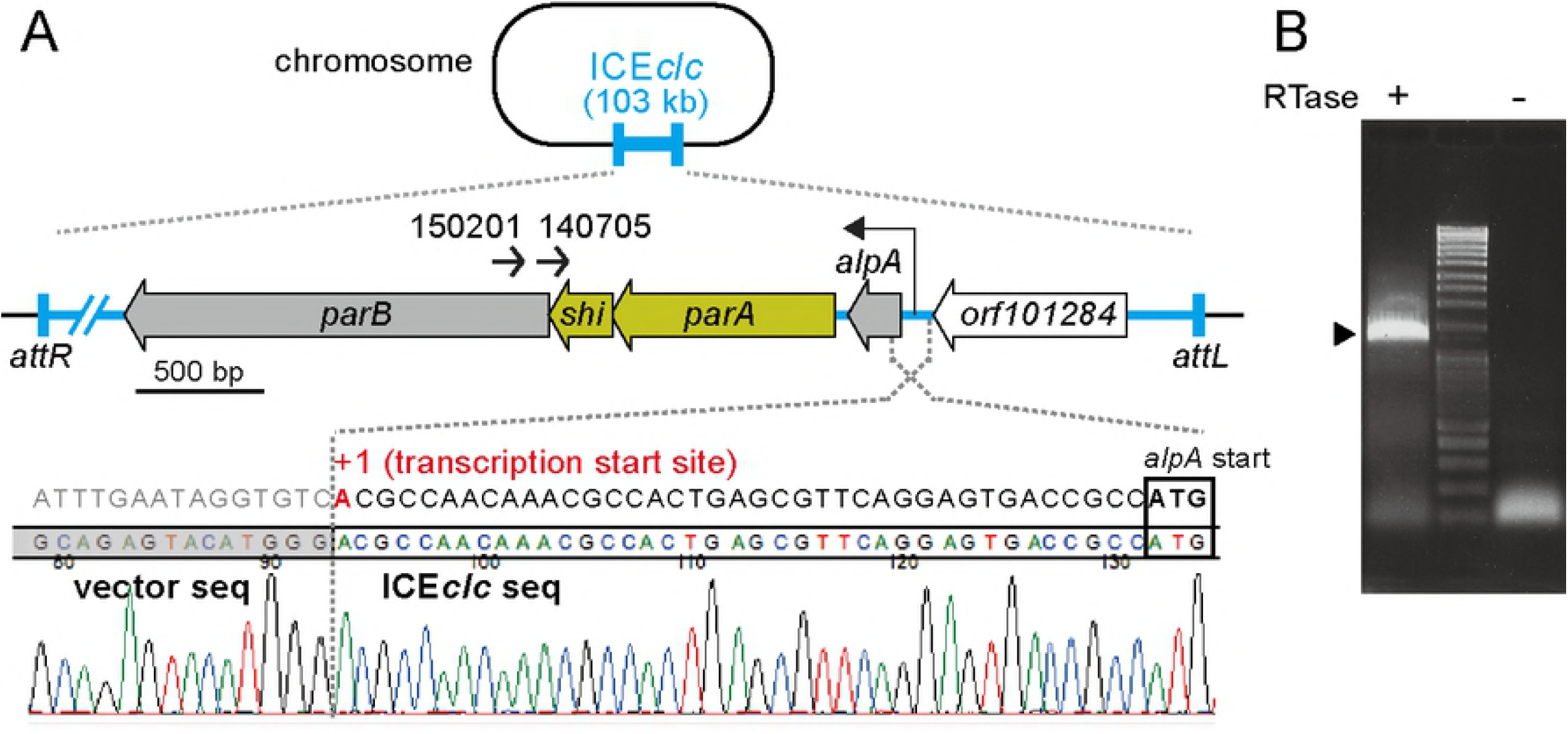
The *parA-shi* region of ICE*clc*. (A) Location of *parA-shi* gene cluster on ICE*clc* nearby the *attL* end. Positions and directions of primers used for 1st (150201) and 2nd (140705) PCRs to amplify 5’RACE products are indicated as small arrows. Lower part shows the sequence of a 1.4-kb 5’RACE product which determined the transcription starting site (adenine in red) of the *parA-shi* cluster. The boundary between the 5’RACE product and the cloning vector used is illustrated. (B) Agarose gel electrophoresis of the 2nd PCR product in 5’RACE. A specific 1.4-kb product is indicated by an arrow head. Presence or absence of the reverse transcriptase in each cDNA synthesis reaction is shown by ‘+’ or ‘-’, respectively.

Here we study genetic characteristics of *parA* and *shi* and their inhibitory effects on host cell growth. We developed fluorescence reporter strains to visualize intracellular behavior of ParA and Shi proteins at the single-cell level and tracak their expression levels as well as host cell proliferation by time-lapse microscopy. Combining experimental and theoretical approaches, we then tested two hypotheses explaining the growth inhibition: division blocking model or elongation blocking model. Our quantitative data and numerical simulations mainly support the former model that the two proteins inhibit cell division, but also indicated that they affect cellular elongation as time elapsed. The results of our study give mechanistic insight into the cellular growth inhibition, which is a highly conserved system in ICE*clc*-related elements.

## Results

### Genetic characterization of *parA-shi* locus

While *parA* has been annotated as encoding a partitioning ATPase, *shi* is predicted to be a short open reading frame (258 bp) of hypothetical protein overlapping 17 bp and 8 bp with up- and downstream genes, *parA* and *parB*, respectively. We thus first examined whether the *shi* gene encodes a protein or acts as a non-coding RNA or a cis-acting element. Two different *shi* mutants which compromised its translation either by the replacement of the start codon ATG with ACG *(shi* mt1) or by the substitution of leucine 7 to an opal mutation *(shi* mt2) were cloned on a vector pME6032 together with *parA* under the control of LacI^q^/P_tac_ system. While the induction of wild-type *shi* with *parA* led to severe growth impairment, the two mutations restored the growth (Fig. 2). This result indicated that *shi* encodes a protein acting with *parA* for the growth inhibition. To determine the transcriptional starting site of the *parA-shi* locus, we performed 5’ rapid amplification of cDNA ends (5’RACE) with reverse primers specifically annealing to the *parB* region. Subcloning and sequencing a specific 1.4-kb 5’RACE product revealed that the transcription started from an adenine 39-base upstream of the *alpA* gene, encoding a putative transcriptional regulator (Fig. 1). This reveals that the gene cluster at least from *alpA* to *parB* are cotranscribed.

**Fig 2.**
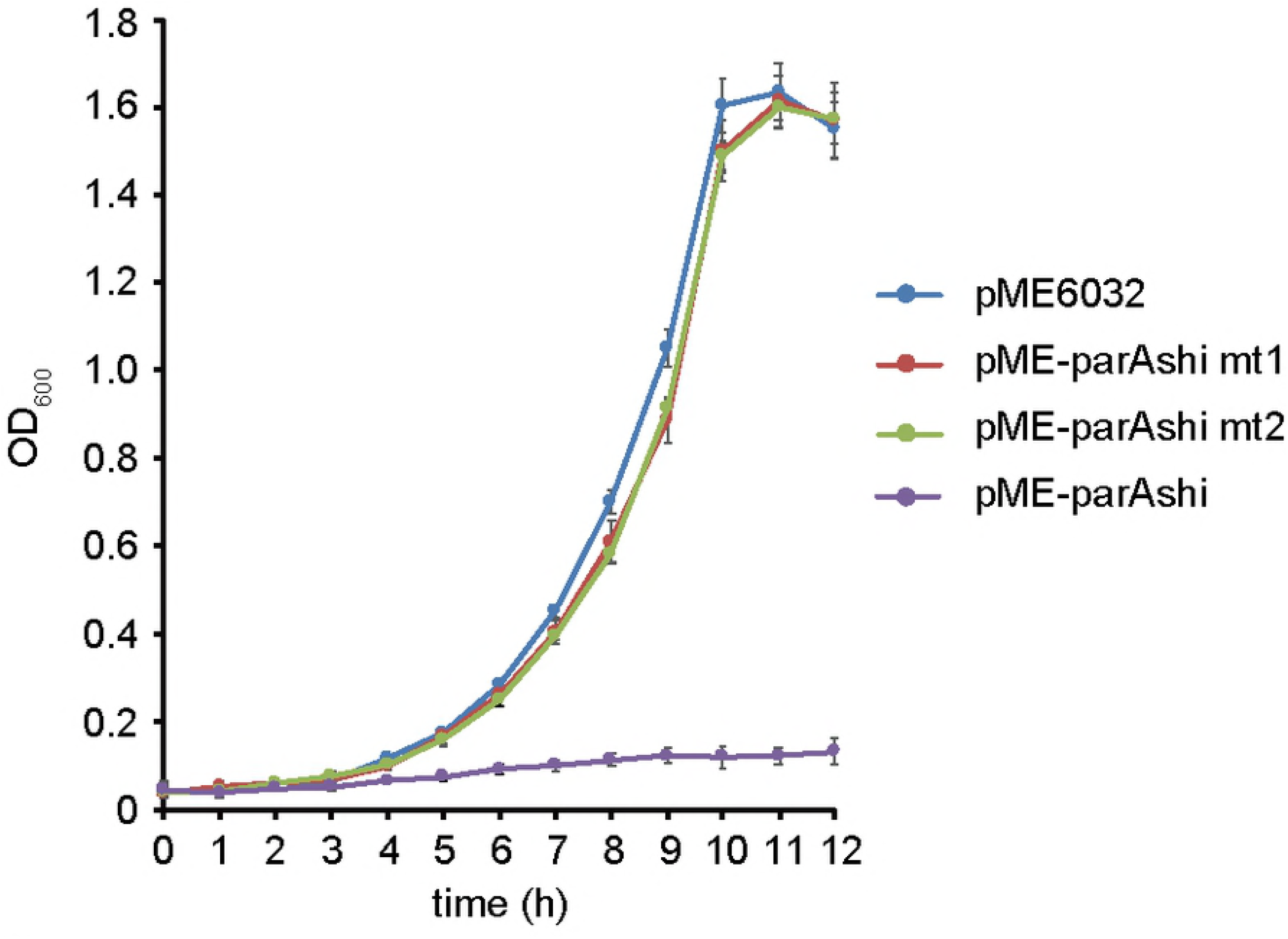
Effect on population growth of different point mutations in *shi*. *P. putida* UWC1 cells carrying pME6032 with different *parA-shi* fragments are cultured with IPTG, and their culture turbidity is measured. Error bars represent standard deviation (SD) from the mean in triplicate assays.

### Phylogeny of ParA and Shi

The *parA-shi* locus is highly conserved in ICE*clc*-related elements found in alpha-, beta-, and gamma-proteobatcterial genomes, except apparently lacking *shi* in ICEHin1056 and SPI-7 (Fig. 3A). ParA of ICE*clc* contains a specific variant of the canonical Walker A motif (KGGxxKT/S), an ATP-binding site, in its N terminal. Phylogenetic analysis of Walker ATPases positioned ParA orthologs encoded on ICEs into a single clade divergent from other ParA-family proteins involved in nucleotide partitioning (Fig. 3B), suggesting that ICE-derived ParA proteins have diverged from other Walker ATPases specific for partitioning chromosomes or plasmids. On the other hand, Shi had been annotated as a hypothetical protein without any domains or motives presented in public databases at a statistically significant level (*E*<0.01). In the current protein database of NCBI, however, we found that some hypothetical proteins that show <80% similarities to Shi contained the HicA_toxin domain (pfam07927, *E*<0.01), a ribonuclease domain conserved in the HicA toxin of the type II toxin-antitoxin (TA) system [19]. Genes of those hypothetical proteins were found in various proteobacterial genomes, such as *Xhanthomonas, Pectobacterium, Pseudomonas*, and *Dickeya*, in the proximity of *parA-* and parB-like genes, while the *hicA* gene of the TA system is adjacent to its cognate *hicB* gene, encoding the anti-toxin protein. Phylogenetic analysis positioned Shi and its relatives into a different clade divergent from HicA toxins (Fig. S1). These phylogeny and conserved syntenies suggest that Shi is diverged from HicA of the type II TA system.

**Fig 3.**
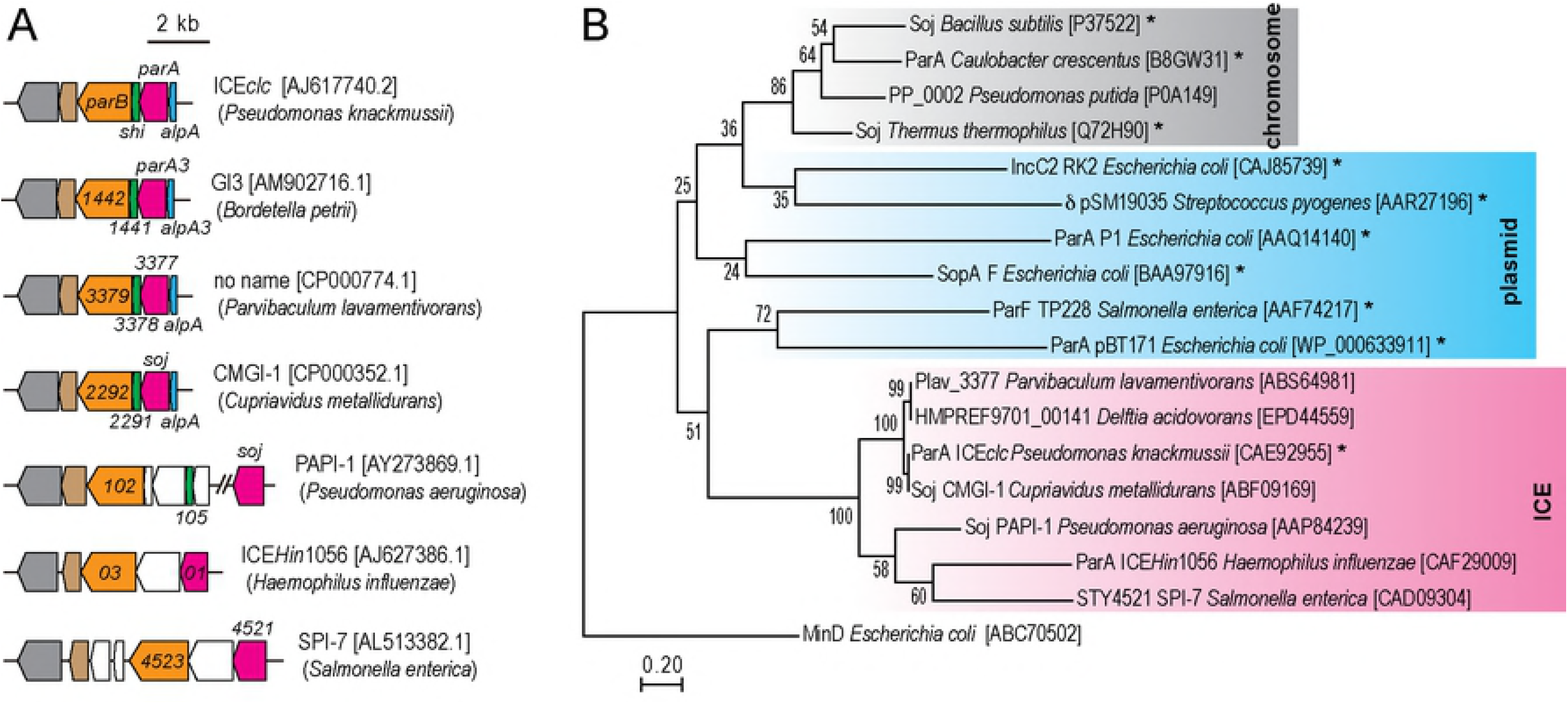
Genetic organization and phylogenetic analysis of ParA proteins. (A) Schematic illustration of *parA-shi* locus on various ICEs. ICE names (if named), species names and accession numbers are shown. Genes are indicated as in the respective genome accession. Common colors indicate similar predicted functions. (B) Maximum-likelihood (ML) tree based on the amino acid sequences of Walker ATPase family proteins. The 18 protein sequences are aligned and used for construction of the ML tree by using the Jones-Taylor-Thornton model. MinD is used as an outgroup. The bootstrap values are shown on each branch. The tree is drawn to scale, with branch lengths measured in the number of substitutions per site. Proteins of which functions are experimentally demonstrated are denoted by asterisks. Accession numbers are shown in brackets.

### ParA-Shi-mediated growth inhibition in heterologous hosts

Given that ICE*clc*-related elements were distributed in a wide range of proteobacterial genomes, we examined the inhibitory effect of ParA-Shi on other proteobacterial cell growth. Induction of ParA and Shi from the plasmid pME6032 significantly inhibited the growth of both *Cupriavidus necator* H16 (beta-proteobacteria) and *E. coli* MG1655 (gamma-proteobacteria), although MG1655 was more susceptible to the inhibitory effect (Fig. 4). Despite repeated attempts to introduce the plasmids with and without *parA-shi* into *Sphingobium japonicum* UT26 (alpha-proteobacteria), no transformants were obtained when the plasmid contained *parA-shi*, probably due to leaky expression of ParA-Shi from the plasmid resulting in inhibition of the colony formation. These results suggest that ParA and Shi still exert inhibitory effect on cell growth of other proteobacteria.

**Fig 4.**
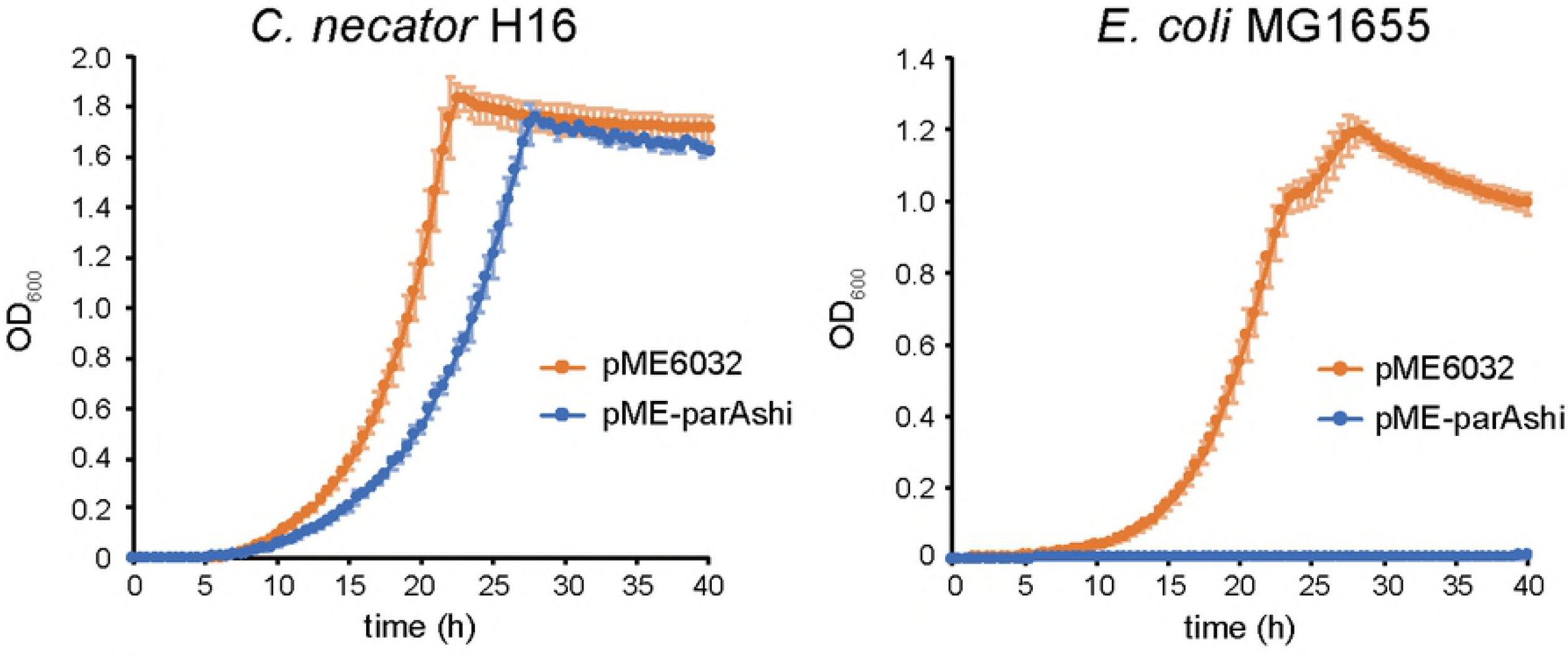
Effect of *parA-shi* expression in different host bacteria. *C. necator* H16 (left) and *E. coli* MG1655 (right) cells carrying either pME6032 or pME-parAshi are cultured with IPTG, and their culture turbidity is measured. Error bars represent SD from the mean in triplicate assays.

### Intracellular localization of ParA and Shi proteins

To investigate mechanistic aspects of the ParA-Shi-mediated cell growth inhibition, we fused ParA and Shi with different fluorescence proteins and observed their intracellular behaviour. Induction of the two fusion proteins, ParA-mCherry and Shi-eGFP, from pME6032 by the addition of IPTG led to growth arrest and elongation of *P. putida* cells (Fig. 5ABC), as observed in the induction of wild-type ParA and Shi (Fig. 2) [14], demonstrating that those fusion proteins still have comparable activities to the wild-type proteins. Standard fluorescence microscopy showed that, within the induced cells, Shi-eGFP apparently diffused in cytoplasm, while ParA-mCherry was colocalized with the fluorescence signal of Hoechst 33342, a dye staining the nucleoid (Fig. 5B). Although the localization of those proteins were retained in cells expressing either ParA-mCherry or Shi-eGFP, growth inhibition and cell elongation were diminished (Fig. 5ABC). These results suggest that ParA binds to chromosomal DNA to disorder cell growth when Shi coexpressed. We then examined whether the association of ParA to chromosome is involved in the cell growth inhibition. Given that the complex formation between Walker ATPase and ATP is required for its binding to DNA [20,21], we generated two ParA-mCherry mutants of which the conformational change was abolished by substituting the N-terminal lysine 15 in the Walker A motif to either glutamine (K15E) or glutamic acid (K15Q). Induction of those mutated proteins with Shi-eGFP did not inhibit cell growth nor elongate cell shape (Fig. 5AD). The different consequences between wild-type and mutant ParA-mCherry were not due to the expression levels of the fluorescence proteins (Fig. 5E). Furthermore, superresolution microscopy using the Airyscan detector revealed that, while wild-type ParA-mCherry colocalized with the nucleoid, ParA(K15E)-mCherry and ParA(K15Q)-mCherry locally accumulated to create aberrant foci (Fig. 5D). The superresolution images also uncovered that Shi-eGFP tends to localize to the cell membrane with multiple foci, irrespective of the ParA mutations (Fig. 5D). These results show that the mutations in ParA abolishing its complex formation with ATP alter subcellular localization of ParA, which results in overriding the inhibitory effect of ParA and Shi on the cell growth.

**Fig 5.**
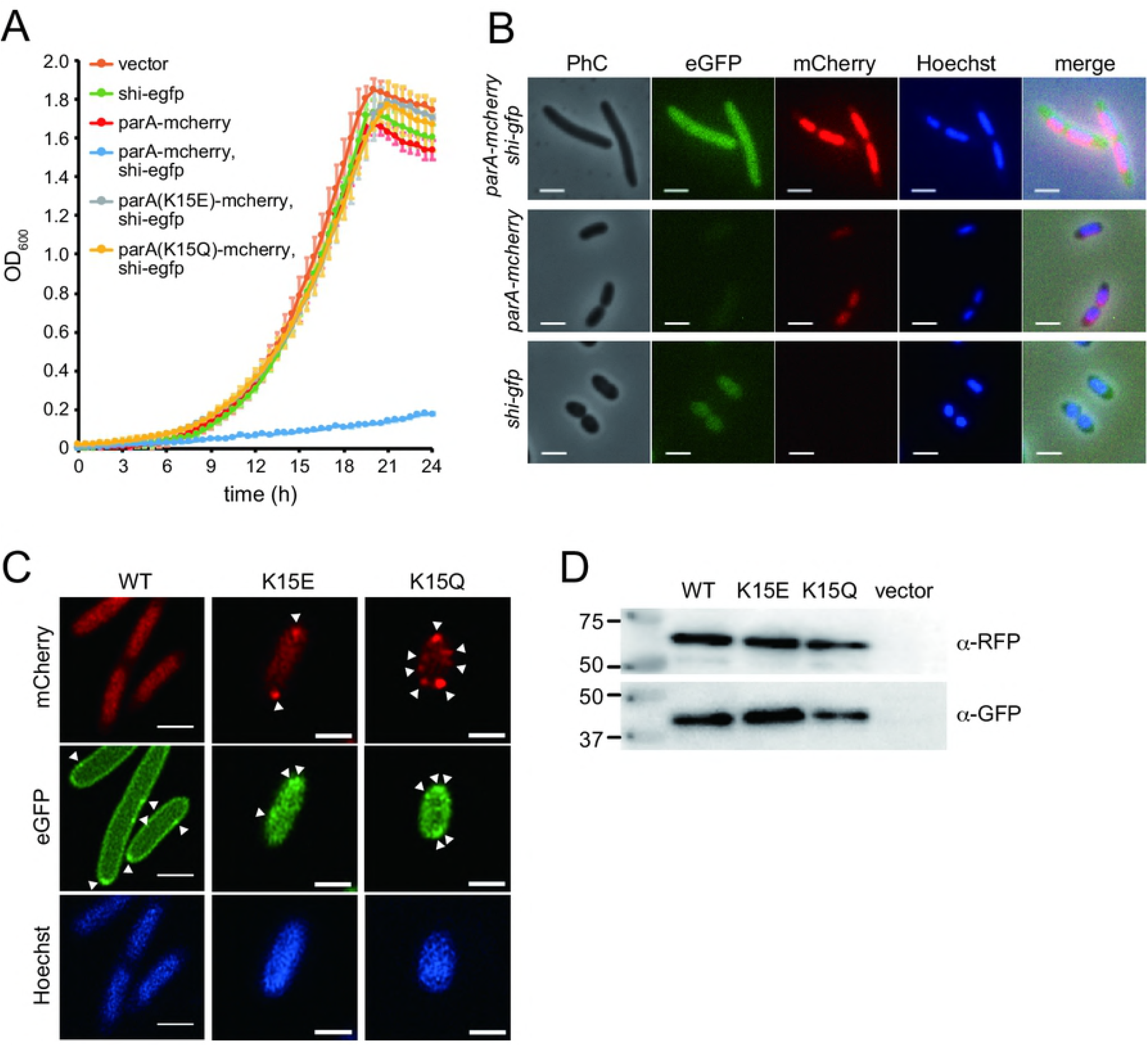
Intracellular localization of ParA and Shi proteins. (A) Effect of fluorescence fusion and site-specific mutations in ParA and Shi on *P. putida* UWC1 population growth. Cells carrying pME6032 derivatives are cultured with IPTG, and their culture turbidity is measured. Genetic information expressed from Ptac promoter on the vector is indicated. Error bars represent standard deviation (SD) from the mean in triplicate assays. Vector, pME6032 (empty). (B) Representative micrographs of *P. putida* UWC1 cells expressing either ParA-mCherry or Shi-eGFP, or both. Phase contrast (PhC) and fluorescence (eGFP, mCherry, and Hoechst33342) images are acquired at 4h after IPTG induction. Scale bar indicates 2 μm. (C) Representative superresolution images of *P. putida* UWC1 cells expressing Shi-eGFP and either ParA(Wild-type)-mCherry, ParA(K15E)-mCherry or ParA(K15Q)-mCherry. Fluorescence images are acquired at 4h after IPTG induction. Aberrant foci are indicated by white triangles. Scale bar indicates 1 μm. (D) Western blot analysis of ParA-mCherry and Shi-eGFP expression in *P. putida* UWC1. Same strains as (C) are cultured with IPTG, and their cell lysates are prepared after 2h induction. Characteristics of ParA (Wild-type, K15E, or K15Q) is indicated in the top, and primary antibodies are in the right. Vector, cells carrying pME6032 (empty).

### Dynamics of cellular growth inhibition by ParA and Shi

To understand dynamics of the inhibitory effect of ParA and Shi on host cell growth, we tracked cellular proliferation and protein expressions at single-cell level using time-lapse microscopy. We used *P. putida* UWC1 expressing ParA-mCherry and Shi-eGFP under the control of LacI^q^/P_lac_ system, and monitored ~200 cells in the presence or absence of IPTG (Movies S1 and S2). Based on the time-course data of the time-lapse observations, we first examined whether the expressions of the two proteins inhibit cellular growth by calculating doubling time of the cells. Induction of the ParA and Shi proteins significantly increased the doubling time of the cells (Fig. 6), indicating that expression of these two proteins decelerate the growth of the cells. We next investigated how ParA and Shi inhibit cellular proliferation. If the two proteins mainly inhibit cellular division process, the IPTG induction would increase cell length but might not affect elongation rate of the cells (Fig. 7A, Hypothesis 1). In contrast, if these proteins prevent cellular elongation process, the rate of elongation would decrease but the cell length would not change by the induction (Fig. 7A, Hypothesis 2). To test these hypotheses, we measured individual cell length and elongation rate in growing microcolonies with or without IPTG. On the beginning of the time course (up to ~200 min), the average cell length in the presence of IPTG was indistinguishable from that in the absence of IPTG (Fig. 7B and S2). As time elapsed, the length was increased by IPTG, whereas that in the absence of IPTG was relatively constant (Fig. 7B and S2). Concomitantly, the maximum cell length, determined as a maximum value of the cell length from birth to division events, was significantly increased by the presence of IPTG, compared to that without IPTG (Fig. 7C). Strong positive correlation between cellular doubling time and maximum cell length was also observed when ParA and Shi expressed, whereas no correlation was observed without induction (Fig. 7D). Thus increasing of maximum cell length and doubling time would be coincident when two proteins were produced. On the other hand, neither elongation rate of cells nor its correlation with doubling time were significantly changed by the induction (Fig. 7EF). These results support the hypothesis that ParA and Shi impair cellular proliferation by inhibiting cellular division rather than elongation process.

**Fig 6.**
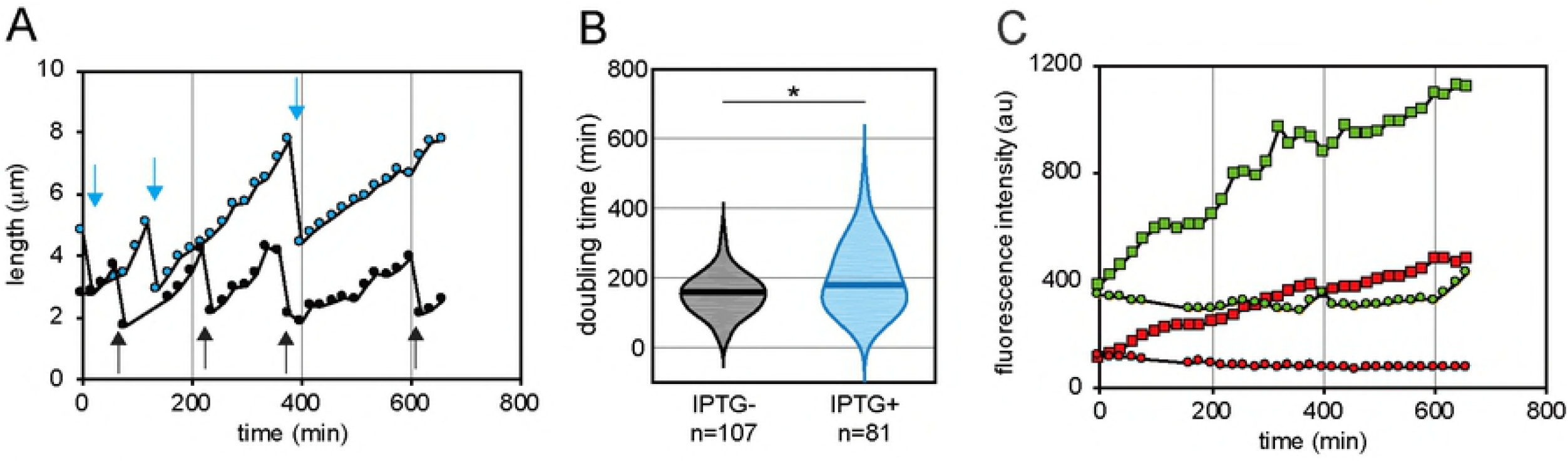
Cellular growth dynamics and changes of fluorescence intensities of ParA and Shi proteins at single-cell level. (A) Temporal changes of cellular length in the absence (black) or presence (blue) of IPTG in a representative cell lineage. Arrows indicate timings of division events. (B) Violin plots of doubling time with or without IPTG induction. Solid lines in plots indicates median of doubling time in each condition. Asterisk indicates significance of difference (P<0.005) in Wilcoxon test. (C) Transitions of fluorescence intensities from Shi-eGFP (green) and ParA-mCherry (red) with (square) or without IPTG induction (circle) in a representative cell lineage. Time indicates the duration from the start of the experiments.

**Fig 7.**
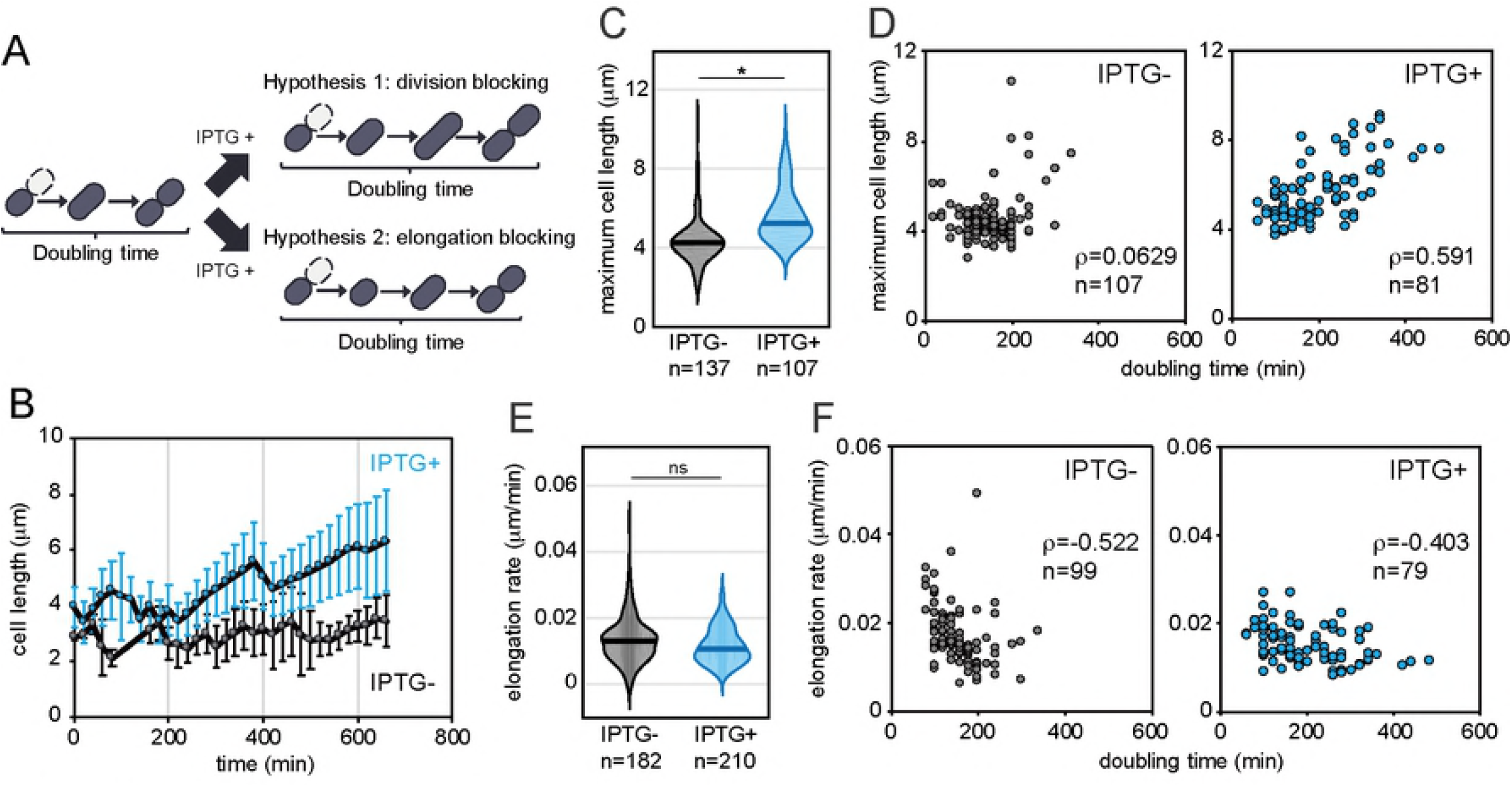
Experimental test of two working hypotheses for ParA-Shi-mediated cell growth inhibition. (A) Schematic illustrations of two scenarios of growth inhibition: division blocking or elongation blocking. (B) Transitions of mean cell length in representative single colonies with (blue circle) or without IPTG (grey circle). Average cell length are calculated from total cells at each time point in each colony. Error bars indicates standard deviations. (C) Violin plots of maximum cell length in the presence and absence of IPTG induction. Cells which did not divide and keep elongation until the end of the experiments are eliminated from analysis. Solid lines indicates median of maximum cell length in each condition. Asterisk indicates significance of difference (P<0.005) in Wilcoxon test. (D) Correlation between maximum cell length and doubling time with (right panel) or without IPTG (left panel). Same datasets as presented in Figure 6B are used. Spearman’s correlation coefficient (ρ) and the number of cells used for analysis (n) are indicated. (E) Violin plots of cellular elongation rate in the presence and absence of IPTG induction. Elongation rate of each cell is calculated by dividing the difference between start and final cell length with elapsed time. Cells which existed less than 4 frames (60 minutes) or did not elongate over 120 minutes are eliminated from analysis. Note that the difference between two conditions is not statistically significant (p>0.005, Wilcoxon test). Solid lines indicates median elongation rate in each condition. (F) Same as (D), but between elongation rate and doubling time.

### Numerical simulation of cell division model

To assess whether the ParA-Shi-mediated inhibition mechanisms suggested by our experiments could adequately elucidate the cellular growth dynamics of *P. putida* cells, we developed a simple cell division model composed of three ordinary differential equations (see Materials and Methods for details). The model consists of three variables: cell length (*L*), amount of cell division machinery (*D*), and concentration of inhibitor proteins, i.e. ParA and Shi, (*P*) (Fig. 8A). Expression of the proteins inhibits elongation and division process and thus causes an increase of cell length and a decrease of elongation rate progressively (Fig. 8B). We also consider the effects of variability among cells independent of the ParA and Shi expressions by defining two parameters *ΔL*_0_ and *Δd*_0_ which denote the maximum rates of elongation and accumulation of division machineries, as probability distribution functions based on our experimental data (see Materials and Methods and Fig. S3 for details). In order to estimate how inhibition of cell division or elongation processes affect cellular proliferation quantitatively, we then performed numerical simulations for hundreds of lineages when the inhibitors have different effects on those two processes (*K_p1_* ≠ *K_p2_*). When inhibitors expressed (*Δp_0_* = 0.01) and mainly stopped cell elongation (*K_p1_* ≪ *K_p2_*), the maximum length of cells did not drastically change, while the doubling time significantly increased (Fig. 8C, left panel; Fig. S4). No correlation between maximum cell length and doubling time was observed even in the presence of inhibitors (Fig. 8C, left panel). Negative correlation between cellular elongation rate and doubling time did not change by the presence of inhibitors (Fig. S5). In the case of division blocking model (*K_p1_* ≫ *K_p2_*), on the contrary, the maximum cell length increased and significantly positively correlated with doubling time, when inhibitor proteins were produced (Fig. 8C, right panel). However, negative correlation between cellular elongation rate and doubling time became weaker by inhibitors production (Fig. S5). These characteristics are consistent to the experimentally observed growth dynamics (Figs 6 and 7), supporting that the growth impairment by the two proteins mainly caused by inhibition of cell division process rather than elongation process.

**Fig 8.**
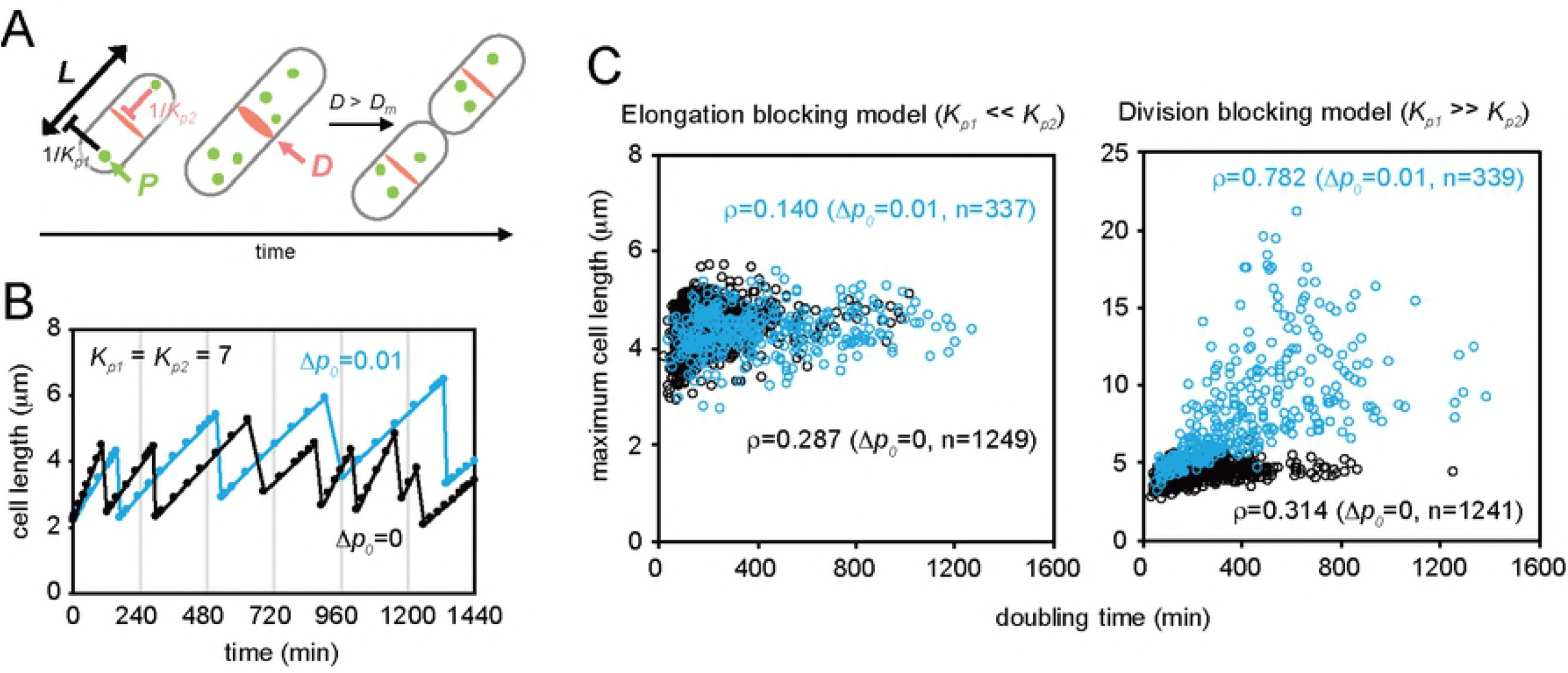
Modeling of cell division. (A) Schematic diagram of a cell division model. The variables are shown in italic boldface. A cell can divide when a sufficient amount of division machineries (*D*) accumulate in a cell (*D* > *D_m_*). Proteins (*P*) are produced at rate Δp_0_ and have negative effects on elongation and/or division processes. Strength of inhibitions on the two processes are controlled by two parameters, *K_p_1* and *K_p2_* (elongation and division, respectively). (B) Examples of numerically simulated growth dynamics in single lineage at different protein production rate (*Δp_0_* = 0 or 0.01). (C) Correlation between maximum cell length and cellular doubling time when different values of *K_p1_* and *K_p2_* are applied to the model (Elongation blocking model, *K_p_1* = 5, *K_p2_* = 200; Division blocking model, *K_p1_* = 200, *K_p2_* = 5). In both conditions, growth of 200 lineages were simulated. Spearman’s correlation coefficient (*ρ*) and the number of cells used for estimating (n) are indicated. The following parameter sets were used for the simulations: *Δp_0_* = 0.01, *D_m_*= 4.4, (*L_0_, P_0_, D_o_*) = (2.2, 0, 2.2).

In both our experimental test and mathematical model, negative correlation between cellular elongation rate and doubling time was observed independent of the inhibitor expression (Fig. 7F and S5), yet no correlation between maximum cell length and doubling time was detected in the absence of inhibitors (Fig. 7D and 8C). We found in the simulation that this interrelation only arose when a variability in elongation rate (CV_elongation_) is larger than that in cell length (CV_length_) (Fig. S6), and our experimental results indeed supports this model: CV_elongation_ (0.710) was larger than CV_length_ (0.289) (Fig. S3), and the correlation coefficient between cellular elongation rate and doubling time (ρ = −0.522) was stronger than that between maximum cell length and doubling time (ρ = 0.0629) (Fig. 7DF). Hence, the strong negative correlation between doubling time and elongation rate is due to larger intrinsic cellular variability in the elongation process.

### Transition of cellular growth inhibition by increasing ParA and Shi expression levels

To investigate relevant expression levels of ParA and Shi for the cellular growth impairment, we quantified their expressions at the single cell level, based on their fused fluorescence proteins. Mean expression levels of ParA and Shi in each cell were negatively correlated to the elongation rate of the cell (Fig. 9A), indicating that the two proteins affect the elongation rate. This observation, however, somehow conflicts with the above result that elongation rate of cells was not significantly changed by ParA-Shi induction (Fig. 7E). Therefore, we hypothesized that the elongation rate would not drastically change as a whole but gradually decrease as ParA and Shi expressions progressively increased with time. The expression levels of ParA and Shi was indeed increased with the birth time of cells (Fig. 9B), indicating that the two proteins were continuously produced after cell division. On the other hand, the cellular elongation rate decreased with the birth time regardless of IPTG induction, but the decline of the elongation rate was more obvious in the presence of IPTG (Fig. 9C). Statistical comparisons of cellular elongation rates revealed that they were indistinguishable between the presence and absence of IPTG in earlier cells (<320 min), whereas the rate with IPTG was significantly lower than that without IPTG in later cells (>320 min) (Fig. 9D).

**Fig 9.**
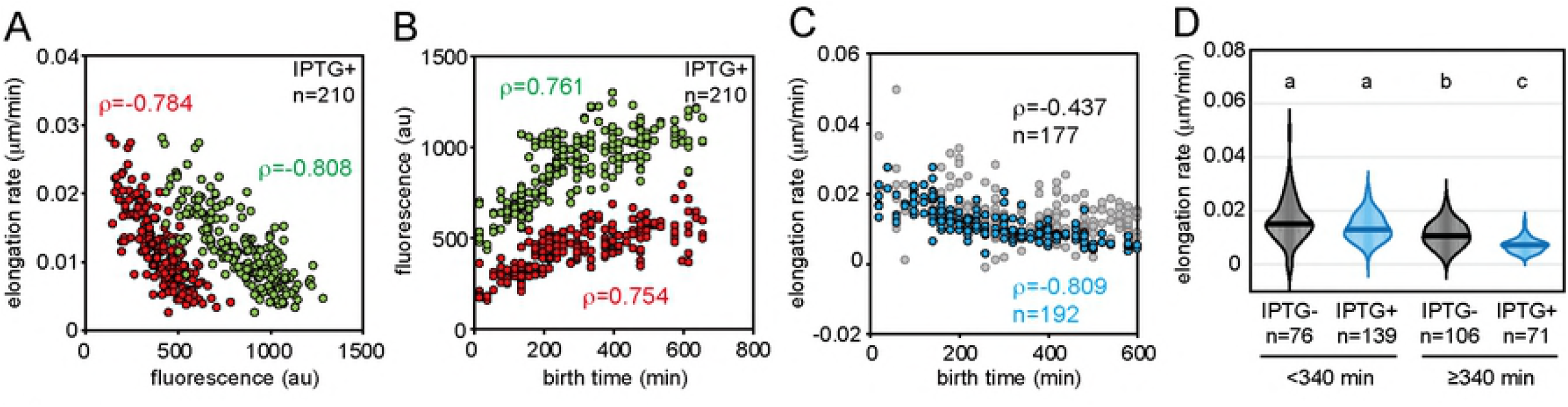
Cellular elongation rate depending on ParA and Shi expression levels. (A) Average elongation rate and ParA-mCherry (red) and Shi-eGFP (green) expression levels in individual cells with IPTG induction. Cells which already existed at the start of the experiments are eliminated from analysis. Spearman’s correlation coefficient (p) and the number of cells used for analysis (n) are indicated. (B) Cellular birth time and average ParA-mCherry (red) and Shi-eGFP (green) expression levels in same cells used in (A). Birth time of an individual cell is defined as the time at which its mother cell divided. (C) Cellular elongation rate and birth time in the presence (blue) or absence (gray) of IPTG. (D) Violin plots of cellular elongation rates in the different birth time with (blue) or without IPTG induction (gray). Solid lines indicate median values in each condition. Letters above plots show significance group based on ANOVA followed by Dwass-Steele-Cirtchlow-Fligner post hoc test (p<0.005).

## Discussion

While mobile DNA vectors often provide selective advantage with host bacteria by transmitting conditionally beneficial functions, such as antibiotic resistance, their presence and activity within the cells can be disadvantageous for the host, since additional physiological and energetic costs may arise from their replication, transcription and translation, as well as horizontal retransfer to the other hosts [22]. To minimize such costs and consequently increase fitness of mobile DNAs in a given ecological niche, they have evolved a variety of system. Some plasmids encode H-NS-like proteins to silence derogative functions that may impede plasmid maintenance [23,24]. Another well-known system in plasmids is the genetic addiction by toxin-antitoxin (TA) genes, through which plasmid-free daughter cells accidentally emerging during cell proliferation are killed by toxin proteins that are more stable than antitoxins, assuming that such daughter cells can become competitors growing faster than cells still carrying plasmids [25]. Temperate phages express regulatory proteins, such as cI repressor in phage lambda, that not only stably maintain the lysogenic state in the host but also control host metabolic pathways to ensure host survival and efficient reproduction of phages [26–28]. As a system for ICEs, we have reported that ICE*clc* invokes a bistable decision between vertical and horizontal transmission [1]. The ICE dedicates a 3-5% subpopulation of host cells (i.e. tc cells) to its costly horizontal transmission, keeping a balance between host and ICE fitness: sacrificing a small proportion of host cells, while maintaining a low but sufficient rate of horizontal transmission of the ICE [17]. Combining experimental and theoretical analyses, we here unveil the dynamics of cellular growth impairment induced in tc cells by ICE-encoding proteins ParA and Shi. Given that the ParA-Shi-mediated growth inhibition results in efficient horizontal transmission of ICE*clc* [14], this system plays a crucial role on successful transfer of the ICE from the limited number of tc cells, by directing them to the costly mission rather than proliferation.

The *parA-shi-parB* locus is highly conserved in other ICE*clc*-like elements found in many different proteobacteria, such as *Acidovorax, Burkholderia, Bordetella*, and *Pseudomonas* species including *P. aeruginosa* [12,13]. ParA and ParB encoded on ICEs show high similarity at amino acid sequence level to those for chromosome or plasmid partitioning (Fig. 3). Nucleotide partition is the most important system for stable segregation of those replicons to daughter cells. A class of the partition systems involves a specific DNA sequence on the segregating replicon that functions as the bacterial equivalent of centromere (e.g. *parS* site), and two proteins: one binds to the centromere (e.g. ParB) and the other is a Walker ATPase with non-specific DNA binding activity (e.g. ParA) [29,30]. Although the mechanism of the partition system with Walker ATPases is still under debate, recent studies more support the diffusion ratchet model in which ParB stimulates ATP hydrolysis of ParA, resulting in the destabilization of ParA-DNA binding and dynamic gradients of ParA-ATP complex as the driver of ParB-DNA cargo [31–33]. Using fluorescence fusion proteins and microscopic imaging, we here showed that ParA of ICE*clc* colocalizes with the chromosome of *P. putida* (Fig. 5). Two amino acid substitutions we made at the ‘signature’ lysine residue of the Walker A motif in ParA, ParA(K15E) and ParA(K15Q), are supposed to interfere with the formation of ParA-ATP complex [21,34]. The former mutation prevents the protein from binding to ATP, whereas the latter still permits the binding but disturb the proper conformational change in the complex. We found that both mutants completely abolished the inhibitory effect on cell growth and the association with DNA (Fig. 5), suggesting that ParA of ICE*clc* also forms a complex with ATP, which is required for the growth inhibition. Considering such functional analogies, one could assume that the competition between ParA proteins from ICE*clc* and chromosome for DNA binding causes the growth impairment. However, the mechanistic details may not be so simple, because ParA of ICE requires Shi but not ParB for the growth inhibition, i.e. the inhibitory effect observed regardless of the presence or absence of ParB [14].

Shi is a curious protein in terms of its action and phylogenetic context. Protein sequences and phylogenetic analysis showed that, although some Shi homologues contain the HicA_toxin domain (pfam07927), they are obviously divergent from HicA toxins of the type II TA system (Fig. S1). HicA is an RNA interferase, causing mRNA and transfer mRNA degradation [35], and thus induction of HicA results in growth impairment of the host [35], whereas Shi did not when expressed alone (Fig. 5A) [14]. Furthermore, our superresolution microscopy revealed that Shi proteins favourably localized in the cell membrane (Fig. 5C). These characteristics suggest that Shi is functionally distinct from HicA toxin, although they might have diverged from a same ancestor.

Our time-lapse imaging and quantitative analysis of cellular proliferation at the single-cell level enable us to draw a several conclusions about the basic principles of growth impairment by ParA and Shi. First, we clearly showed that ParA and Shi impair the growth of *P. putida* by blocking cellular division rather than elongation (Fig. 7). We then developed a simple model to explain the cellular proliferation of free-living bacteria (Fig. 8, see Materials and Methods). Our numerical simulation using the mathematical model rationally shows that the division blocking model proposed in our experiments is feasible to explain the growth dynamics of *P. putida* cells with IPTG induction: increase of doubling time and cell length by ParA and Shi expression result from their disturbance of cellular division process rather than elongation process (Fig. 8). Intriguingly, we also found that the intrinsic variance in elongation rate is generally larger than that in cell length among *P. putida* cells (Fig. S3). Although it is too early to conclude that this observation is common in any free-living bacteria, it suggests that the cellular length or the timing of cell division are more strictly controlled than the elongation rate in this species. Moreover, it is noteworthy that growth dynamics and inhibitory effect obviously change by increasing intracellular levels of ParA and Shi. We found that, although the cell division is the main process blocked by ParA and Shi, the elongation is gradually, yet not drastically as an entire period, interfered by ParA and Shi expression (Fig. 9). Further analysis is still needed to determine whether this change is caused by direct actions of ParA and Shi or an indirect consequence of other cellular functions.

In conclusion, our results provide qualitative and quantitative aspects of cell growth inhibition by ParA and Shi proteins as an adaptive strategy of ICE*clc* for increasing its horizontal transfer frequency and fitness. The mathematical model we developed here can be simple but, with further knowledge of host-ICE partnerships, become more versatile for describing and predicting dynamics of microbial populations in which ICEs horizontally transfer.

## Materials and Methods

### Bacterial strains and culture media

*Escherichia coli* DH5α (Gibco Life Technologies) and DH5αλpir [17] for plasmid constructions was routinely grown at 37°C on LB medium [36]. *E. coli* MG1655 [37], *Cupriavidus necator* H16 [38], and *Pseudomonas putida* UWC1 [39] were cultured at 30°C on LB or type 21C minimal medium (MM) [40] containing either 10 mM Fructose or 5 mM 3-chlorobenzoate (3CBA). If necessary, antibiotics were added at the following concentrations; kanamycin 25 μg mL^−1^, gentamicin 20 μg mL^−1^, ampicillin 100 μg mL^−1^, and tetracycline 10 μg mL^−1^ for *E. coli*, 25 μg mL^−1^ for *C. necator* and 50 μg mL^−1^ for *P. putida*.

### DNA techniques and plasmid constructions

Preparation of plasmid and chromosomal DNAs, digestion with restriction endonucleases, DNA fragment recovery, DNA ligation, and transformation of *E. coli* cells were carried out according to established procedures [36] or to specific recommendations by the suppliers of the molecular biology reagents (Qiagen and Takara). The transformation of bacterial cells by electroporation was performed as described previously [41]. Routine PCR was performed with ExTaq or PrimeStar DNA polymerase (Takara), and primers used are listed in Table S1. All PCR products cloned were confirmed by sequencing with the BigDye Terminator version 3.1 (Applied Biosystems) and an ABI PRISM 3700 sequencer (Applied Biosystems). Plasmids used in this study are listed in Table S2. To introduce point mutations in *shi*, inverse PCR was carried out using pME-parAshi as a template with two different primer sets (140701 and 140702, or 140703 and 140704). Each PCR product was self-ligated, transformed in *E. coli*, and verified for correctness of the *parA-shi* locus with point mutations. To avoid PCR-based errors on the vector part, the 1.1-kb fragment containing *parA* and mutated *shi* genes was recovered from each plasmid by EcoRI-XhoI digestion, and recloned on a fresh pME6032. To produce a C-terminal fusion of ParA to mCherry (i.e. ParA-mCherry), a ~900 bp fragment containing the *parA* gene without its stop codon was amplified using pME-parAshi as a template and primers (140801 and 140802). The fragment was cloned in EcoRI and *HindIII* sites on pBAM-link-mcherry [Miyazak PLosGen], resulting in pBAM-parA-link-mcherry. A ~1.7-kb fragment including the *parA-mcherry* fusion gene was obtained by EcoRI-SpeI digestion of pBAM-parA-link-mcherry, subcloned into the same sites on pBluescriptIISK+, and then recovered by EcoRI-SacI digestion. The fragment was cloned in the same sites on pME6032 to generate pME-parA-mcherry. To produce a C-terminal fusion of Shi to eGFP (i.e. Shi-eGFP), we first amplified a ~750 bp fragment containing the *egfp* gene using pJAMA23 [42] as a template and primers (140301 and 140302), in which the start codon of *egfp* was replaced by a short nucleotide sequence encoding 15 amino acids (KLPENSNVTRHRSAT) as a linker peptide. The fragment was cloned in *HindIII* and SpeI sites on pBAM-link-mcherry, resulting in pBAM-link-egfp. A ~280 bp fragment containing the *shi* gene without its stop codon was then amplified using pME-parAshi as a template and primers (140804 and 140805). The fragment was cloned in *Eco*RI and *Hind*III sites on pBAM-link-egfp, resulting in pBAM-shi-link-egfp. A ~1-kb of *EcoRI-Bgl*II fragment from pBAM-shi-link-egfp was cloned into EcoRI-BglII sites of pME6032 to generate pME-shi-egfp. The ~1.7-kb of EcoRI-SpeI fragment from pBAM-parA-link-mcherry and a ~1kb of *XbaI-Bgl*II fragment from pBAM-shi-link-egfp were together cloned in the *EcoRI-Bgl*II sites on pME6032 for generating pME-parA-mcherry-shi-egfp.

### 5’ rapid amplification of cDNA ends (5’RACE)

Total RNA was isolated from *P. putida* clc6 cells grown in MM with 5 mM 3CBA until stationary phase, by using RNAprotect Bacteria Reagent and RNeasy Mini kit (Qiagen), following manufacture’s instruction. To remove contaminating genomic DNA, an 8 μg of the isolated RNA was further treated with 2U of TURBO DNase (Invitorgen) at 37°C for 1 h and purified with RNeasy spin columns (Qiagen). The amount of RNA was quantified with Qubit RNA BR assay kit (Invitrogen). A 500 ng of the RNA was used for the 5’RACE reaction with SMARTer RACE 5’/3’ kit (Clontech), according to manufacture’s instruction. In brief, a first-strand cDNA was synthesized by SMARTScribe Reverse Transcriptase, SMARTer II A oligonucleotide, and a specific primer 150201 that anneals the 5’ region of the *parB* gene. Using the cDNA as a template, the 1st 5’RACE PCR was performed with primers 150201 and Universal Primer Mix (UPM) provided with the kit. To increase specificity, the 2nd PCR was carried out with the 1st PCR product as a template using primers 140705 and short UPM provided with the kit. The 2nd PCR product (1.4 kb) was purified from an agarose gel and cloned into the provided pRACE vector. The plasmid carrying the 5’RACE product was sequenced with M13.R primer to determine the transcription starting site of *parA* and *shi* genes.

### Phylogenetic analysis

The amino acid sequences of Walker ATPases involved in either chromosome or plasmid partitioning and those orthologous encoded on ICEs were obtained from the NCBI GenBank. The sequences were aligned with the program MUSCLE (https://www.ebi.ac.uk/Tools/msa/muscle/), and the Maximum-likelihood (ML) tree was then reconstructed using MEGA 7.0.26 with Jones-Taylor-Thornton model, Nearest-Neighbor-Interchange (NNI) move and 100 bootstrap replicates [43]. A discrete Gamma distribution was used to model evolutionary rate differences among sites. All positions containing gaps and missing data were eliminated. There were a total of 185 positions in the final dataset. The tree was rooted with MinD from *E. coli* MG1655. Alignment of the amino acid sequences of Shi from ICEs with close hits in the GenBank nr database and construction of the tree were performed with the same procedure described above. The tree was generated using a total of 49 positions and rooted with MazF from *E. coli* MG1655.

### Bacterial growth test in liquid media

Bacterial strains were pregrown in MM with fructose and tetracycline until stationary phase and adjusted to OD_600_=3.0. The preculture was reinoculated with 0.1% dilution into the fresh medium containing 1 mM IPTG. Optical density (OD_600_) was measured every 0.5 h, and its mean and standard deviation were calculated by biological triplicates.

### Microscopy

*P. putida* UWC1 derivatives were precultured in MM with fructose until stationary phase and then diluted 1% into the fresh medium containing 1 mM IPTG. After 4 h incubation, cells were stained by Hoechst33342. Cells and fluorescent proteins were imaged with a Zeiss Axio Observer epifluorescence microscope (Carl Zeiss). Images were taken with a Axiocam 506 monochrome camera (Carl Zeiss), a 100x/1.40 oil immersion Plan-Apochromat lens (Carl Zeiss) at exposure times of 350 ms for phase contrast and 100 ms for fluorescence images. The light source and filter used for fluorescence imaging was Zeiss Colibri7 and Filter Set 81 HE, respectively. Images were digitally recorded as 16-bit TIFF-files using the Zeiss Zen software, and analyzed using METAMORPH (Molecular Devices). Super-resolution images were observed using a LSM800 confocal laser scanning microscope, equipped with a 100x/1.46 oil immersion alpha Plan-Apochromat lens and an Airyscan detector (Carl Zeiss). mCherry, eGFP, and Hoechst33342 were excited with 561 nm, 488 nm, and 405nm lasers, respectively. Airyscan processing was performed with the 2D SR mode. Images for display were artificially colored ‘red’ (for mCherry), ‘green’ (for eGFP), or ‘blue’ (for Hoechst 33342), and then auto-leveled and cropped to the final resolution and image size using Adobe Photoshop (Adobe Inc.).

### Time-lapse experiments

*P. putida* cells containing the pME-parA-mcherry-shi-egfp plasmid were pre-cultured in LB medium supplemented with tetracycline for over 16 h. The culture was diluted to the density of OD_600_=0.04 with MM plus 10 mM fructose and tetracycline. A 10 μL of the diluted culture was plated onto 1.0 % agarose pad of the MM in the closed cultivation chamber (H. Saur Laborbedarf). The chamber was connected to a syringe by 1×2 mm silicone tubes, and filled with the liquid MM with or without 0.2 mM IPTG by CX07100 syringe pump (Isis, Japan) on the beginning of the experiments.

Time-lapse measurements of the cellular growth in the closed cultivation chamber were performed using a microscope, Axio Observer with ZEN software and a motorized stage unit (Carl Zeiss). Stabilization of Z-offset in each position were facilitated by the use of Definite Focus (Carl Zeiss) during the experiments. The cultivation chamber and the microscope were kept at 30°C in an acrylic box with heater unit (Tokken, Japan) during the time-lapse experiment. Images were acquired with an Axiocam 506 mono and a 100/1.40 oil lens with Colibri LED excitation light source (Carl Zeiss). The cells were exposed for 100 ms using 475nm and 555nm, both at 15% power, for taking eGFP and mCherry fluorescence images, respectively. For the acquisition of phase contrast images, transillumination LED light was irradiated to the cells for 100 ms at 3.8 V power.

The time-lapse interval was 20 min. For the data analysis, we used the images acquired after 60 min after the start of the experiments to wait for the temperature and the culture condition in the chamber to be stable on the beginning of the experiments.

### Image processing and parameterization of cellular characteristics

Five (IPTG-) and 11 micro-colonies (IPTG+) on the agarose pad were randomly chosen for analysis. Firstly, we processed the microscopy images using ImageJ (NIH) for subtracting background signals and enhancing the signals from the region of the cells. Based on these processed images, we made binary mask images by detecting cell contours, using Schnitzcells [44], a MATLAB based software (MathWorks, USA). In IPTG-conditions, we used the processed phase-contrast images for making masks. For generating the mask images in IPTG+ conditions, we used eGFP fluorescence images because the eGFP fluorescence distributed in cytoplasm region and facilitated clear detection of cell boundaries. Using these mask images, we tracked individual cell across multiple images by Schnitzcells. Finally, we obtained the several information for individual cell such as duration, transition of cell length, division (birth) time, change of eGFP and mCherry mean fluorescence intensity and daughter-parent relation. These data were further processed as necessary.

### Western blot analysis

*P. putida* UWC1 derivatives carrying pME6032, pME-parA-mcherry-shi-egfp, pME-parA(K15E)mcherry-shi-egfp, or pME-parA(K15Q)mcherry-shi-egfp were precultured in MM with fructose until stationary phase and then diluted 1% into the fresh medium containing 1 mM IPTG. After 2 h incubation, cell lysates were subsequently extracted by sonication, and their protein concentrations were measured using Protein Assay kit (BIO-RAD). We subjected the lysates (2 mg of protein per sample) to 12.5% SDS-PAGE analysis and immunoblotting. Rabbit polyclonal antibody to GFP (1:1000, MBL) and rabbit polyclonal antibody to RFP (1:1000, MBL), were used as primary antibodies, whereas horseradish peroxidase-conjugated antibody to rabbit (1:1000, Cell Signaling) was used as secondary antibodies. Immobilon Western (Millipore) was used for detection. Images were captured with ChemidocTM XRS+ systems (BIO-RAD).

### Model descriptions

Our cell division model consists of three ordinary differential equations as follows.

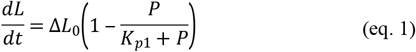

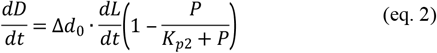

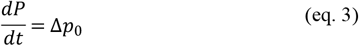

In bacteria, linear and exponential growth models are most commonly used [45], and both models could sufficiently depict the experimentally observed growth curves in the range of our time-lapse experiments. For simplicity, we here assumed that a cell length (*L*) extends linearly at constant rate, *ΔL_0_*, (eq. 1) based on the time-lapse observations (Fig. 7). In broad range of the prokaryotes and unicellular eukaryote, a cellular division event is thought to closely couple to their length or size [46,47], and thus a cell should extend to a proper length threshold before its division event by monitoring its size or increment from birth [48]. For simplicity, here we assume that the rate of synthesis or formation of division machinery (*dD/dt*) is proportional to the rate of cell extension (*dL/dt*) (eq. 2), and the cell divides into two progenies when the certain quantity of the division machineries accumulate in individual cell (*D* > *D_m_*). After the division, a cell body and division machineries are thought to be equally distributed, and thus the length and the amount of division machinery becomes half of the original quantity in the model (*L_t+1_* = 1/2 *L_t_, D_t+1_* = 1/2 *D_t_*) (Fig. 8A). We also introduced the concentration of the proteins (i.e. ParA and Shi) as variable *P* (eq. 3), and its negative effect on both elongation and division processes (eq. 1 and 3). Given that the concentration of the both two proteins almost constantly increased in the presence of IPTG (Fig. 6C), the protein production rate can be assumed to be constant (denoted as *Δp_0_*) in the model. We also could suppose that the concentration of those proteins is not affected by cell division (i.e. *P_t+1_ = P_t_ + Δp_0_*). In addition, we found that there were originally heterogeneous characteristics in the cellular elongation rate and the division timing (i.e. cell length before the division) independent of the expressions of the proteins (Fig. 6 and 7, and Fig. S3), and we defined the two parameters, *ΔL_0_* and *Δd_0_*, as probability distribution functions based on the quantitated experimental data from time-lapse measurements (see Fig. S3).

## Acknowledgements

We thank Hitomi Matsuo, Kayo Ohkouchi, Mutsumi Sakurai for their technical assistance.

## Supporting Information

**Fig S1. Maximum-likelihood (ML) tree based on the amino acid sequences of HicA and Shi-like hypothetical proteins.** The 19 protein sequences are aligned and used for construction of the ML tree by using the Jones-Taylor-Thornton model. MazF, another type II toxin, is used as an outgroup. The bootstrap values are shown on each branch. The tree is drawn to scale, with branch lengths measured in the number of substitutions per site. Proteins which contain the HicA_toxin domain (pfam07927, *E*<0.01) are denoted by asterisks.

**Fig S2. Example of transitions of mean cell length in single colonies with or without IPTG induction.** At each time point, average cell length are calculated for total cells in the colony. Error bars indicates standard deviations. Colony i1-i3, with IPTG; Colony n1-n3, without IPTG.

**Fig S3. Distributions of maximum cell length and elongation rate in the absence of IPTG induction.** Histograms and distributions of the maximum cell length and the rate of cell elongation in the absence of IPTG were shown. The data were fitted to Burr distribution using MATLAB fitdist function, and the obtained parameter sets were used for the generating probability distribution of *ΔL_0_* and *Δd_0_*. Because we wished to estimate the variability of growth parameters in normal cells, we excluded the data of non-elongating cells, which stopped elongation over 2 hours, from this fitting.

**Fig S4. Violin plots of the cellular doubling time in numerical simulation using elongation or division blocking models.** In both cases, induction of protein inhibitors (*Δp_0_* = 0.01) significantly increased cellular doubling time (p<0.005, Wilcoxon test). Solid lines indicates median of doubling time. Used parameter sets are same as described in Fig. 8.

**Fig S5. Scatter plots of cellular elongation rate and doubling time obtained by numerical simulations.** Elongation blocking (left) and division blocking (right) models are shown. Spearman’s correlation coefficient (ρ) and the number of cells used for analysis (n) are indicated. Used parameter sets are same as described in Fig. 8.

**Fig S6. Innate variability in cellular elongation and division processes affects correlation between doubling time and elongation rate or maximum cell length.** Plots of absolute values of Spearman’s correlation of the indicated pairs (correlation between doubling time and elongation rate, red; or maximum cell length, blue) when CV of elongation rate and maximum cell length changed. In this simulation, we set *ΔL_0_* and *Δd_0_* as functions of normal distribution and changed their variance from 0.01 to 0.5 and from 2 ×10^−4^ to 1×10^−2^, respectively. For each parameter set, growth of 200 lineages were simulated, and the obtained results were used for calculations of ρ and CV. The following parameter sets were used for the simulations: *Δp_0_* = 0, *D_m_*= 4.4, (*L_0_, P_0_, D_0_*) = (2.2, 0, 2.2). The results of numerical simulations without inhibitor production (*Δp_0_* = 0) showed that the strong negative correlation between elongation rate and doubling time only arose when a coefficient of variation of innate cell elongation rate (CV_elongation_) is larger than that of maximum cell length (CV_length_). In the opposite way, where CV_elongation_ is smaller than CV_length_, maximum cell length rather than elongation rate strongly correlated to the doubling time.

**Movie S1. Time-lapse movie of *P. putida* UWC1 carrying pME-parA-mcherry-shi-egfp, cultured on agarose-pad without IPTG induction.**

**Movie S2. Time-lapse movie of *P. putida* UWC1 carrying pME-parA-mcherry-shi-egfp, cultured on agarose-pad with IPTG.**

**Table S1. Oligonucleotides used for PCR amplification.**

**Table S2. Plasmids used in this study.**

